# EEG functional connectivity metrics wPLI and wSMI account for d distinct types of brain functional interactions

**DOI:** 10.1101/450270

**Authors:** Laura Sophie Imperatori, Monica Betta, Luca Cecchetti, André Canales Johnson, Emiliano Ricciardi, Francesca Siclari, Pietro Pietrini, Srivas Chennu, Giulio Bernardi

## Abstract

Functional connectivity (FC) estimation methods are extensively used in neuroimaging to measure brain inter-regional interactions. The weighted Phase Lag Index (wPLI) and the weighted Symbolic Mutual Information (wSMI) represent relatively robust exemplars of spectral (wPLI) and information-theoretic (wSMI) connectivity measures that recently gained increased popularity due to their relative immunity to volume conduction. wPLI and wSMI are posited to have different sensitivity to linear and nonlinear relationships between neural sources, but their performance has never been directly compared. Here, using simulated high-density (hd-)EEG data, we evaluated the accuracy of these two metrics for detecting distinct types of regional interdependencies characterised by different combinations of linear and nonlinear components. Our results demonstrate that while wPLI performs generally better at detecting functional couplings presenting a mixture of linear and nonlinear interdependencies, only wSMI is able to detect exclusively nonlinear interaction dynamics. To evaluate the potential impact of these differences on real experimental data, we computed wPLI and wSMI connectivity in hd-EEG recordings of 12 healthy adults obtained in wakefulness and deep (N3-)sleep. While both wPLI and wSMI revealed a relative decrease in alpha-connectivity during sleep relative to wakefulness, only wSMI identified a relative increase in theta-connectivity, while wPLI detected an increase in delta-connectivity, likely reflecting the occurrence of traveling slow waves. Overall, our findings indicate that wPLI and wSMI provide distinct but complementary information about functional brain connectivity, and that their combined use could advance our knowledge of neural interactions underlying different behavioural states.

## Introduction

Functional connectivity (FC) metrics identify statistical (undirected) associations among spatially distinct brain areas. Electroencephalography (EEG) and magnetoencephalography (MEG) represent popular neuroimaging modalities for the estimation of FC owing to their high temporal resolution, in the order of milliseconds. However, both EEG and MEG suffer from volume conduction, which results from the instantaneous propagation of electric fields generated by a primary current source to all (or most) of the on-scalp sensors. Because of this linear mixing of different sources on the same sensor, common methods for FC estimation, such as coherence or mutual information, may lead to the identification of apparent functional couplings that do not reflect true brain inter-regional interactions [1]–[3]. To overcome this problem, several new FC methods have been specifically designed to minimize the impact of volume conduction effects. In particular, the weighted Phase Lag Index (wPLI, [1]) and the weighted Symbolic Mutual Information (wSMI, [4]), represent examples of spectral (wPLI) and information-theoretic (wSMI) connectivity estimation methods that are increasingly applied to both EEG and MEG data [5]–[14]. These two connectivity metrics are modified versions of pre-existing methods (PLI [1], [15]; SMI [4]) that minimise the contribution of zero-lag interactions potentially determined by volume conduction. These approaches are thus expected to allow identifying true time-lagged functional couplings [16]–[20] in the activity of underlying brain sources, while excluding apparent zero-lag connectivity driven by a mixture of real and spurious relationships [21], [22].

Both wPLI and wSMI have been applied to explore brain functional dynamics associated with different behavioural states [6], [7] or potential network-level alterations in pathological conditions (e.g., Alzheimer’s disease [8], schizophrenia [9] and social anxiety disorder [14]). Interestingly, they have also been suggested to allow the identification of variations in functional integration, accompanying changes in the level of consciousness, following severe brain injury or under anesthesia [4], [10]–[13], [23], [24]. For instance, King and colleagues [4] found that wSMI connectivity between centro-posterior areas and other brain regions is higher in healthy conscious individuals as compared to patients with *unresponsive wakefulness syndrome* (UWS) or in a *minimally conscious state* (MCS). Similarly, Chennu and colleagues [10], [24] showed that alpha-band wPLI-based functional networks differ between healthy individuals and patients with disorders of consciousness (UWS, MCS). In line with this, previous studies [13], [23] also showed that propofol sedation in healthy individuals is associated with a decrease in alpha-band wPLI [23] and a relative increase in delta-band wPLI connectivity [13].

In spite of these promising findings, it is currently unclear whether the two methods provide a similar description of brain inter-regional relationships, or account instead for distinct types of functional interactions. In fact, wPLI [1] is a measure of phase synchronisation that may account for linear interactions but is also expected to be sensitive to nonlinear couplings [25], [26]. On the contrary, wSMI [4] is thought to reveal nonlinear relationships due to its grounding in information theory [27]. However, the actual performance of the two methods at detecting distinct types of connectivity dynamics has never been directly compared in simulated or real experimental data.

Therefore, here we used simulated high-density (hd-)EEG data to specifically investigate and compare the accuracy of wPLI and wSMI in identifying different types of interaction dynamics, including both linear and nonlinear dependencies. In addition, to evaluate the potential impact of differences between the two methods on the analysis of real experimental data, we tested wPLI and wSMI on hd-EEG recordings collected from human participants in distinct behavioural states, namely wakefulness and deep (N3-)sleep, typically characterised by markedly different levels of consciousness [28]. In light of previous observations suggesting that the two methods may allow the detection of differences in the level of consciousness [4], [10], [12], [23], we expected both wPLI and wSMI connectivity to differ between wakefulness and N3-sleep. However, here we also asked whether the two connectivity metrics provide overlapping or complementary information about changes in brain functional dynamics across the two vigilance states.

## Materials and Methods

### Simulation of hd-EEG data

The MATLAB-based (The MathWorks, Inc., Natick, Massachusetts, USA) *’Berlin Brain Connectivity Benchmark’* (BBCB) framework [29] was used to simulate realistic hd-EEG recordings (108 channels, 500Hz, 120s). In particular, the simulated electrical activity was generated by imposing bivariate relationships between two cortical sources, which were then projected at scalp level using a biophysically realistic model of electrical current propagation in the head. The adopted model was based on the standard ICBM152 anatomical template [30] and included 6 tissue types: scalp, skull, air cavities, gray matter, white matter and cerebrospinal fluid (CSF). We modeled both intra and inter-hemispheric interactions between pairs of cortical sources (Figure 1). Specifically, the first source was placed either in left (LIPL) or right (RIPL) inferior parietal lobule, while the second source was kept in the right middle frontal gyrus (RMFG). The choice of these locations was motivated by previous neuroimaging studies showing that resting state activity of these areas is modulated by the level and content of consciousness [31]-[34]. For the sake of simplicity, only two interacting sources at a time were considered: LIPL-RMFG (inter-hemispheric) and RIPL-RMFG (intra-hemispheric).

**Figure 1.**
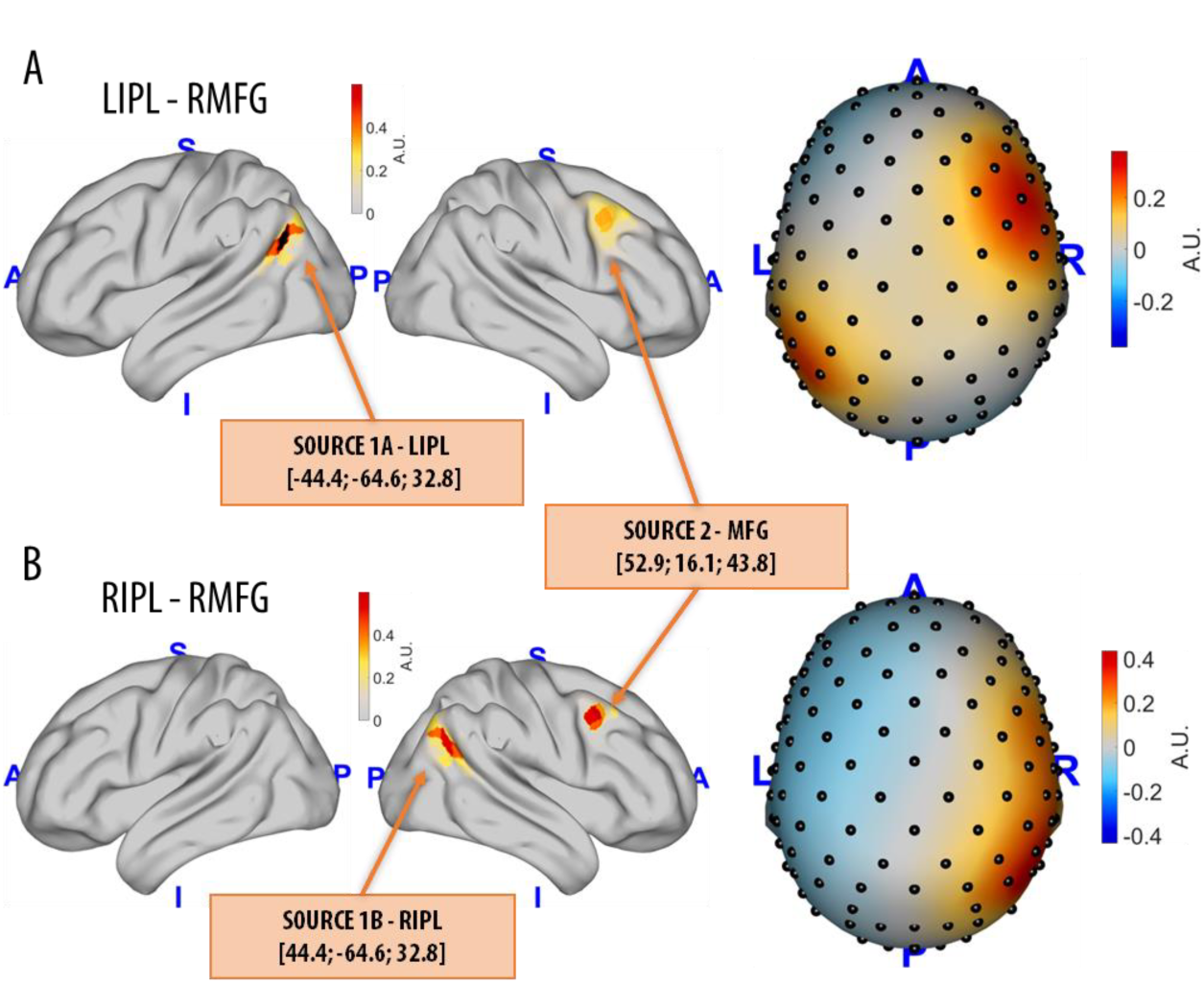
From modelling source dynamics to EEG field patterns. Intra and inter-hemispheric interactions between two source pairs were modelled: the first source was placed either in left (LIPL) or right (RIPL) inferior parietal lobule, while the second source was kept in the right middle frontal gyrus (RMFG). Source amplitudes are shown using a lateral view of the brain, while resulting EEG field potentials are plotted using a top view of the scalp (A.U. stands for arbitrary unit).

As detailed below, we simulated nine different coupling relationships between the two sources, which differed in the type and relative degree of linear and nonlinear components. For each pair of source locations (LIPL-RMFG and RIPL-RMFG) and each type of simulated source coupling dynamics we also modelled 100 different signal-to-noise ratios (SNR; from 0.01 to 1, with steps of 0.01), which describe the weighting of simulated source signals with respect to simulated background activity. As detailed below, 100 different background noise patterns were obtained for each considered SNR. Specifically, brain noise *n*_*b*_(*t*) was generated by placing 500 mutually statistically independent time-series characterised by 1/f-shaped power (pink noise) and random phase spectra at an equal number of random locations sampled from the entire cortical surface. Moreover, spatially and temporally uncorrelated sensor noise *n*_*s*_(*t*) was sampled from a univariate standard normal distribution. The overall noise contribution was defined as noise n(t):

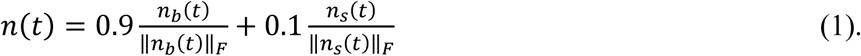

### Source Interaction Dynamics

For each source pairing (LIPL-RMFG, RIPL-RMFG), nine different coupling relationships were simulated by modelling the time-series of the two sources based on linear (AR) and nonlinear (Hènon [35], Ikeda [36], Rössler [37], Lorenz [38]) dynamical systems. *Linear interactions.* The time courses of the two sources were modelled using bivariate linear autoregressive (AR) models of order 5:

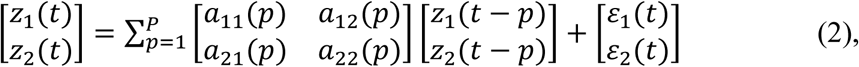

where *a*_*ij*_(*p*), *i,j*∈{1,2}, *p*∈{1,‥,*P*} are linear AR coefficients, and ɛ*i*(*t*), *i*∈{1,2} are uncorrelated standard normal distributed noise variables. The off-diagonal entry *a*_12_(*p*) was set to zero, while *a*_21_(*p*) was set to 0.5. Thus, interactions arise from a unidirectional time-delayed influence of *z*_1_on*z*_2_. Moreover, the generated time series were bandpass-filtered in the alpha band (8-12 Hz) using an acausal third-order Butterworth filter with zero phase-delay [29]. We decided to simulate alpha oscillations with a clearly defined sender-receiver relationship, as they are also a key feature of brain activity in physiological wakefulness [39].

### Nonlinear interactions

The time courses of the two sources were modelled by considering each one as a time-varying state variable of a specific dynamical system. In particular, we considered four different nonlinear systems: two defined by two-dimensional non-iterated maps (Hénon [35], [40] and Ikeda [36]) and two represented by three-dimensional nonlinear ordinary differential equation systems (Rössler [37], [40] and Lorenz [38]). Dynamical systems describe the motion of a point in a multidimensional state space, where the starting point is defined by the initial conditions of the system. For each system all potential combinations of variables have been considered as representing different interaction dynamics, i.e. Hénon (x,y), Ikeda (x,y), Rössler (x,y), Rössler (x,z), Rössler (y,z), Lorenz (x,y), Lorenz (x,z), Lorenz (y,z). The MATLAB-based Chaotic Systems Toolbox was used to compute the time series for the selected nonlinear systems, and the respective parameters were chosen to achieve complex chaotic behaviour: Hénon map [a=1.4; b=0.3], Ikeda map [μ=0.9], Rössler dynamics [a=0.2, b=0.2, c=5.7,*x*_0_=0.1,*y*_0_ =0.1,*z*_0_=0.1,*h*=0.1], Lorenz dynamics [σ=10, β=28, **ϱ**=8/3,*x*_0_=0.1,*y*_0_=0.1,*z*_0_=0.1,*h*=0.1]. Due to the complex nature of these dynamics, they have not been limited to a specific frequency band.

### Connectivity Analysis

The simulated EEG datasets (108 channels, 500Hz, 120 s) generated for each coupling model were divided into 60 non-overlapping 2 s-epochs ([1], [4], [41]). Then, FC was computed for each epoch and pair of electrodes. Analyses were focused on the 0.5-12 Hz frequency range. Before computing connectivity measures, a current-source-density transform [42] was applied to the EEG data, as in previous works [1], [4]. This method provides a reference-independent signal and acts as a spatial filter, leading to a relatively improved spatial resolution [43].*wPLI.* The wPLI measures the extent to which phase angle differences between two time series *x*(*t*) and *y*(*t*) are distributed towards positive or negative parts of the imaginary axis in the complex plane (similar to the PLI [1], [15]). The underlying idea is that only real non-volume conducted activity will give rise to a distribution of phase angles, predominantly on the positive or negative side. The PLI is defined as the absolute value of the sum of the signs of the imaginary part of the complex cross-spectral density*S*_*xy*_ of two real-valued signals *x*(*t*) and *y*(*t*) at time point or trial *t.*

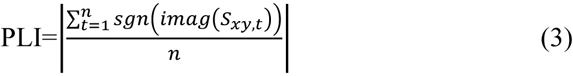

The weighted phase-lag index [1] is an extension of PLI, in which contributions of angle differences are scaled according to their distance from the real axis to address potential further volume conduction confounds:

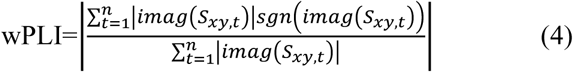

The wPLI is based only on the imaginary component of the cross-spectrum, and thus implies robustness to noise compared to coherence, as uncorrelated noise sources cause an increase in signal power [16]. Here wPLI was computed using the Fieldtrip toolbox [44] (multi-taper method fast Fourier transform, single Hanning taper [1], 0.5 Hz frequency resolution). The mean value across frequency-bins in the 0.5-12 Hz range was computed to obtain a single wPLI coupling value. *wSMI.* The wSMI [4] evaluates the extent to which two EEG signals present non-random joint fluctuations, suggesting sharing of information. The time series *X* and *Y* in all EEG channels are first transformed into sequences of discrete symbols 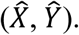 The symbols are coded according to the trends in amplitudes of a specific predefined number of consecutive time points. We chose the kernel *k* to be 3, implying that the symbols are constituted of three elements, leading to 3!=6 different potential symbols in total [4], [11]. The temporal separation of elements that constitute a symbol was set to be τ=14 frames (*τ*_*t*_=*ms*) such that the maximum resolved frequency was 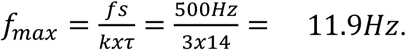

The joint probability of each pair of symbols co-occurring in two different time series is computed to estimate the symbolic mutual information (SMI) shared across two signals. To address volume conduction artifacts, the weighted symbolic mutual information disregards co-occurrences of identical or opposite-sign signals.

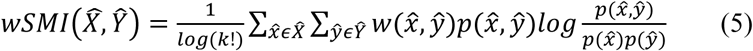

The wSMI can lead to negative values, given that it is a weighted mutual information measure, a form of weighted relative entropy [45].

### Statistical Procedure for Simulated Data

The accuracy of wPLI and wSMI was evaluated at *whole-brain* and *topographic* levels, respectively indicating *i*) the ability to detect the presence of statistical dependencies in the overall (median) connectivity across all pairs of electrodes (see Figure 2), and *ii*) the ability to detect a significant interaction between the pairs of electrodes spatially closest to the actual brain sources among all pairs of electrodes.

**Figure 2.**
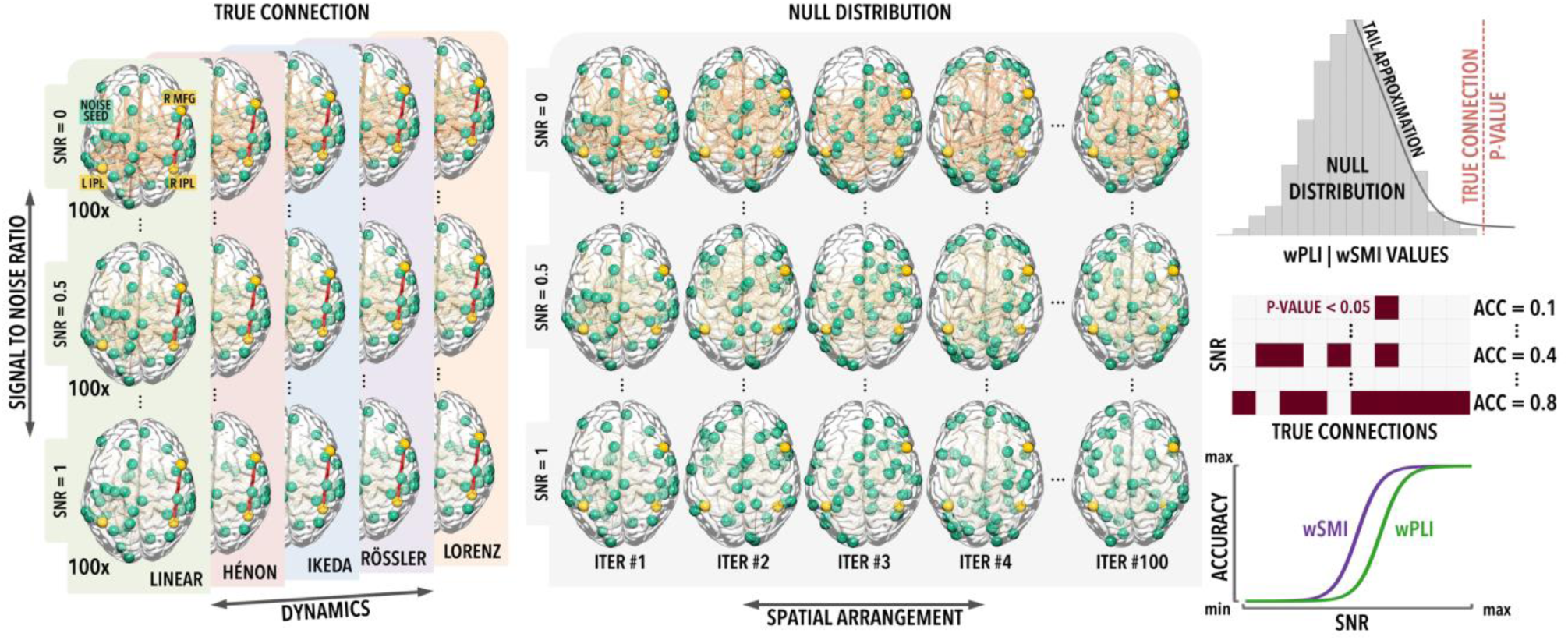
Outline of the methodological design for the assessment of whole-brain accuracy. The source locations LIPL, RIPL and RMFG are marked as yellow dots in the brain plots, while the red line indicates a true interaction between two of the sources (RIPL-RMFG). For each SNR in the range 0.01-1.00 (0.01 steps; *N*=100) different spatial distributions (*N*=100) of random background noise (marked as green dots) were generated in combination with true interactions between the source pairs and projected at scalp level.The corresponding null distributions were obtained through time-point-shuffling of the original interacting source-level timeseries. The same procedure has been applied to all interaction dynamics and tested source pairs (intra/inter-hemispheric).

#### Whole-brain accuracy

For each source pairing (LIPL-RMFG, RIPL-RMFG), interaction dynamics and SNR, the whole-brain detection accuracy of wPLI and wSMI was computed as the proportion of cases (N=100 datasets differing by their respective spatial noise distributions), in which the whole-brain median connectivity value (across all electrode-pairs) of simulated EEG data passed the 95^th^ percentile of a null distribution obtained after time-point-shuffling of the original source-level timeseries (N=100 permutations; Figure 2). To account for the small number of permutations, a generalised Pareto distribution was used to model the tail of the null distribution, using the PALM (Permutation Analysis of Linear Models) software [46]. Of note, we chose to focus on a time-point-shuffling procedure instead of phase-shuffling, since the latter can introduce spurious interdependences between time-series, especially for the Rössler dynamics [47]. However, in the Supplementary Material, we also present results obtained with null distributions generated by phase-shuffling the original time-series using the Amplitude-Adjusted-Fourier-Transform (AAFT) procedure [48], [49]. With the expected exception of the Rössler dynamics, the two approaches provided similar results (see Figure S1).

#### Topographic accuracy

For each source location pairing, interaction dynamics and SNR, the topographic accuracy was defined as the proportion of simulated EEG datasets (N=100 differing by their respective spatial noise distributions), in which the connectivity between the two electrodes spatially closest to the cortical sources (minimum Euclidean distance) passed the 95^th^ percentile of all other electrode pairings (in the same simulated EEG-recording; 5778 channel pairs).

In summary, for both approaches, a threshold corresponding to the 95**^th^** percentile of the respective null-distributions (surrogate data for whole-brain connectivity, and connectivity of all electrode-pairs in topographic analysis) was regarded as the limit for the detection of significant FC interactions (α = 0.05). The mean total accuracy of wPLI and wSMI was computed as the mean of accuracies obtained across all SNRs. Non-parametric permutation tests (N=10000, p < 0.05) were used to compare the performance of the two metrics at each SNR and for mean accuracy. Specifically, for each examined condition, the difference in mean accuracy between wPLI and wSMI was compared with a null distribution obtained by randomly ‘reassigning’ to the two metrics the values of accuracy determined for the different SNR configurations. A similar procedure was used to compare performance of wPLI and wSMI for different spatial distributions of noise at each SNR.

## Experimental hd-EEG recordings

In order to verify whether potential differences between wPLI-and wSMI-based FC measures in recognizing distinct interaction dynamics have actual implications for real-data analysis, an additional investigation was performed on hd-EEG recordings (257 channels, Electrical Geodeisics Inc.; 500 Hz, ~2min) obtained in different behavioural states. Specifically, data were obtained from 12 healthy volunteers (25 ± 4 yrs, 6F) during distinct states of vigilance: relaxed wakefulness with eyes closed (W) and deep (N3-)sleep. The data was recorded as part of a larger project aimed at exploring the effects of changes in visual experiences during wakefulness on NREM-sleep features [50]. Brain activity during N3-sleep was extracted from an overnight EEG recording in the sleep laboratory, whereas wake data consisted of six minutes of eyes-closed resting-state activity obtained at 8AM the following morning, when homeostatic sleep pressure is expected to be at its minimum [51]. Sleep scoring was performed using standard procedures [52] and all 30s epochs containing N3-sleep were extracted and concatenated. All wake and N3-sleep recordings were bandpass filtered between 0.5 and 45 Hz (NetStation 5, EGI) and visually inspected to identify and reject bad channels (which were later interpolated using spherical splines).

EEG recordings during wakefulness were divided into non-overlapping 5s segments and visually inspected to identify and reject clear artifacts. A procedure based on Independent Component Analysis (ICA) was used to remove residual ocular, muscular, and cardiac artifacts [53]. For each subject, we randomly extracted and analyzed the minimum common number (across subjects) of artifact-free 2s-long epochs, corresponding to 70 segments (i.e. 140s; the first 0.5s and the last 0.5s of each 5s segment were discarded). The same amount of data (i.e. 70 2s-epochs; 140s) was randomly selected from N3-sleep that occurred during the first half of the night. From this selection, we excluded epochs representing potential outliers in terms of signal power within classical frequency bands. Specifically, the Power Spectral Density (PSD; Welch’s method, Hamming windows, 8 sections, 50% overlap) of all N3 2s-epochs was calculated in delta (0.5-4Hz), theta (4-8Hz), alpha (8-12Hz), sigma (12-16Hz), beta (18-25Hz), gamma (30-45Hz) and broadband (0.5-45Hz) frequency ranges. Then, outlier segments for any of the seven considered frequency ranges (i.e., threshold = median PSD ± 2 median absolute deviations (MAD)) were excluded from the random selection procedure (see Figure S2). For each condition and channel, the median wPLI and wSMI connectivity of each electrode to all other scalp electrodes was computed in all epochs for the 0.5-12 Hz frequency range (i.e., as in simulated data). The median one-to-all connectivity of each electrode was computed and compared to the average of the median one-to-all connectivity across surrogate datasets (1000 iterations) generated through time-point shuffling of the original recordings of each channel. In this approach, the same permutation scheme was used for all subjects, and the global signal, corresponding to the average signal across all electrodes, was re-added to each shuffled dataset. Paired comparisons were performed between i) wakefulness and surrogate data, and ii) wakefulness and N3-sleep (non-parametric permutation test; p<0.05). Correction for multiple comparisons was ensured using a permutation-based supra threshold cluster correction [54], [55]. In brief, the same contrast was repeated (N=10000 iterations) after shuffling the labels of the two compared sets and the maximum size of significant electrode-clusters was saved in a frequency table. A cluster-size threshold corresponding to the 95th percentile of the obtained distribution (α = 0.05) was applied to correct for multiple comparisons. Whole-brain connectivity (median of one-to-all connectivity across all electrodes) was also evaluated and compared to surrogate data using a non-parametric permutation test (N=10000 iterations; p < 0.05).

## Results

### Linear and nonlinear interdependencies in simulated data

First, we quantified the content of linear and nonlinear interdependencies in the nine examined interaction dynamics: linear AR model, Hénon map, Ikeda map, Rössler (x,y), Rössler (x,z), Rössler (y,z), Lorenz (x,y), Lorenz (x,z), Lorenz (y,z). In particular, to quantify the linear content of the bivariate relationships between the original sources we used cross-correlation, which offers a simple measure of similarity of two signals as a function of the displacement of one relative to the other [56]. In order to measure the nonlinear content, we took the average of the directional, nonlinear interdependence measure N in both directions of the source dynamics [56], [57]. As shown in Figure 3, most of the interaction dynamics we modelled presented a mixture of linear and nonlinear dependencies, with the notable exception of Lorenz (x,z) and Lorenz (y,z), which showed a clear predominance of nonlinear interactions.

**Figure 3.**
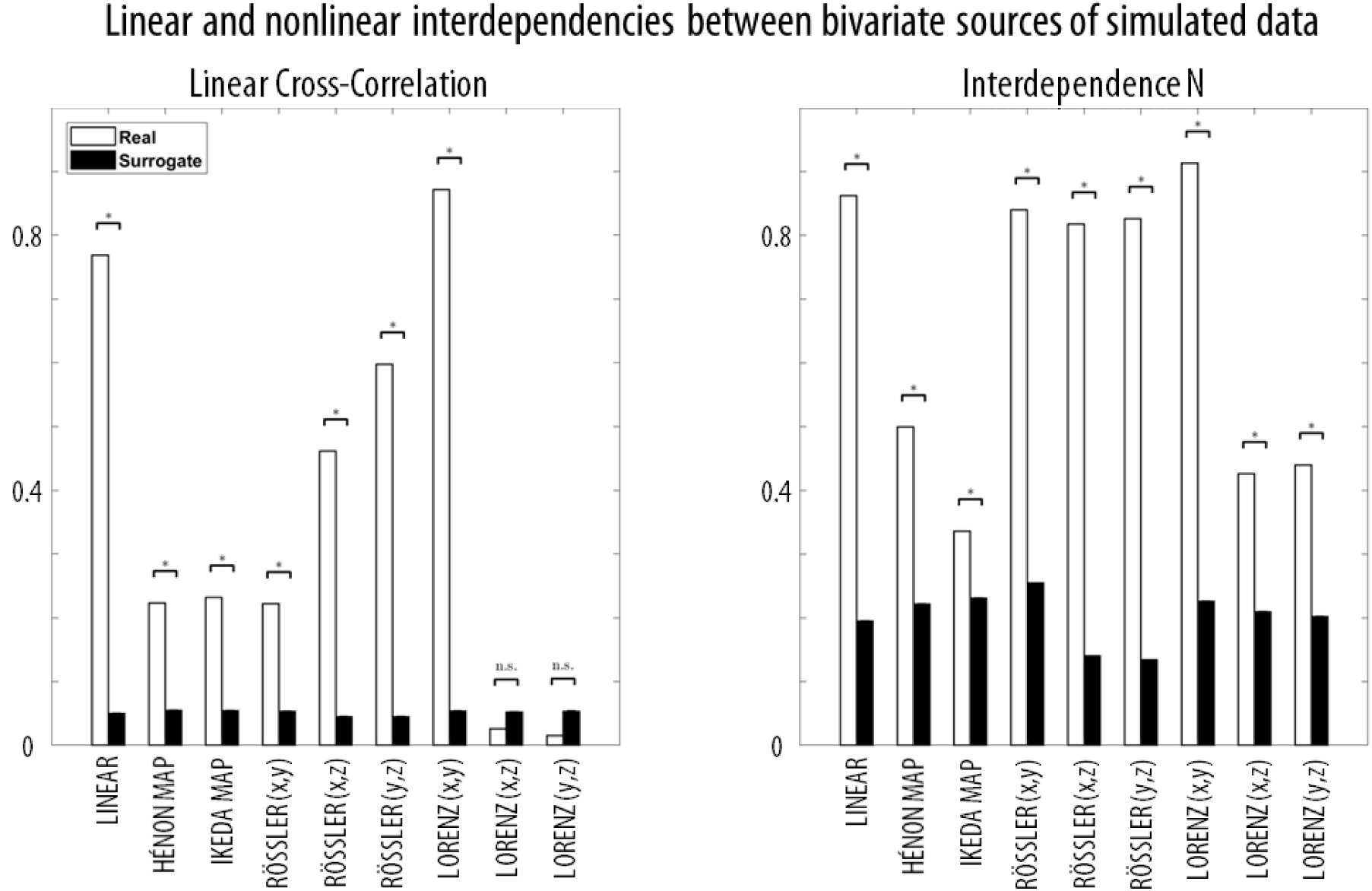
The absolute value of cross-correlation (CC; measure of similarity of two series as a function of the displacement of one relative to the other) and the interdependence measure N (measure of the nonlinear relationship between two time series) were computed for simulated true source time-series (0.5-12 Hz) and the null distribution, obtained by shuflling the shuffled source time-series (N=1000, 0.5-12 Hz). For both CC and N, zero (0) indicates total independence, while one (1) indicates strong dependence. The differences between the true simulated data and its null distribution, i.e. surrogate data, were computed (* for p_one-tail_ < 0.05, Bonferroni-corrected based on 18 comparisons). The error bars show the standard error of the mean for the null distribution. For all cases, N was computed using the following parameters (but very similar results were obtained when optimal, individual parameters were selected for each time-series): embedding dimension (m=10), time lag (tau=5), theiler correction (theiler=50), number of nearest neighbours (nn=10).

### Simulated data - whole-brain connectivity

The whole-brain detection accuracy was computed as the proportion of cases in which the whole-brain median connectivity value of each simulated EEG dataset passed the 95th percentile of a null distribution obtained after time-point-shuffling of the original timeseries. Figure 4A shows the mean accuracy of wPLI and wSMI (averaged over all SNRs) computed for each source pairing (intra/inter-hemispheric) and tested interaction dynamics. Figure 4B shows the whole-brain accuracy at each SNR. Of note, the accuracy of the two connectivity measures was similar for intra and inter-hemispheric connections. The performance of both metrics was similar for the linear relationship in the broadband (0.5-12 Hz) signal. However, wPLI showed higher accuracy than wSMI in the intra-hemispheric case when connectivity in the alpha-band (8-12 Hz; corresponding to the range in which the interaction was modelled), was specifically considered (Figure S3). Moreover, wPLI performed better at detecting Hénon and Ikeda dynamics, especially at high SNRs (≥0.97 and 0.5-0.95, respectively). Both wPLI and wSMI showed significant and comparable levels of accuracy for all Rössler (x,y; x,z; y,z) cases at all tested SNRs, with the exception of low SNRs (Rössler (x,z) SNRs 0.1-0.15; Rössler (x,z) SNRs 0.1-0.2), for which wSMI tended to achieve a better detection performance. For the Lorenz (x,y) dynamics, wPLI achieved a better mean accuracy relative to wSMI, with the strongest differences observed for low SNRs (0.18-0.53). On the other hand, wSMI had higher accuracy for identifying Lorenz (y,z) dynamics for all SNRs ≥0.14. Finally, while no overall performance differences were observed at detecting Lorenz (x,z)-based interaction dynamics, wSMI tended to achieve a higher accuracy for low/intermediate SNRs, between 0.23 and 0.68. Of note, with the expected exception of the Rössler dynamics, similar results were obtained when the null distributions were generated using phase-shuffling (AAFT) instead of time-point shuffling (Figure S1).

**Figure 4.**
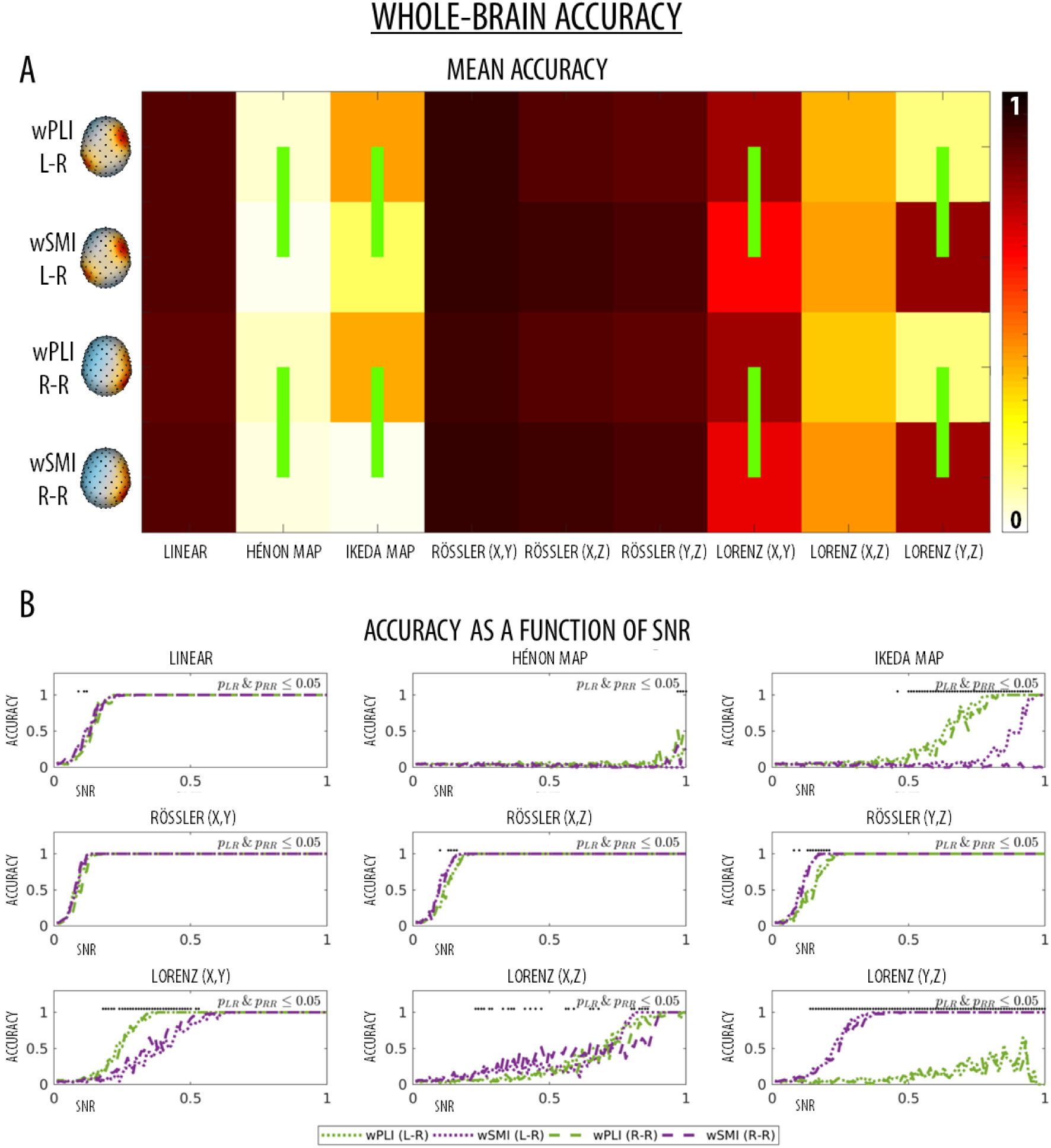
**A)** Mean whole-brain detection accuracy for all nine different relationships between the chosen source location pairings (L = left IPL to right MFG; R = right IPL to right MFG). The green vertical lines mark significant differences between wPLI and wSMI (permutation tests, p < 0.05) for each type of interaction, pairing of source locations and SNR. **B)** Whole-brain detection accuracy for all nine different relationships between the chosen source location pairings as a function of SNRs (L = left IPL to right MFG; R = right IPL to right MFG). Black dots at the top of each graph mark significant accuracy differences between wPLI and wSMI for each specific SNR that were observed for both intra and inter-hemispheric conditions.

### Simulated data - topographic connectivity

The topographic accuracy was defined as the proportion of simulated EEG datasets in which the connectivity between the two electrodes closest to the cortical sources passed the 95th percentile of all other electrode pairings. Results are similar to those described for whole brain accuracy (Figure 5). For the linear dynamics, wPLI and wSMI showed again similar mean accuracies, but wPLI tended to show higher accuracy for low SNRs (0.05-0.09) and high SNRs (> 0.94). Accuracy of wPLI (but not of wSMI) further improved for band-limited connectivity in the alpha-range (8-12 Hz; Figure S3), especially for low SNRs (0.04-0.08) as well as high SNRs (≥ 0.87). For both Hénon and Ikeda iterated maps, the mean topographic accuracy of wPLI was significantly higher than the mean topographic accuracy of wSMI. Specifically, in the Hénon case, wPLI had higher accuracy especially for SNRs ≥ 0.64, while in the Ikeda case, it had higher accuracy at intermediate SNRs (0.31-0.74). Both wPLI and wSMI showed significant and comparable levels of mean accuracy for the three Rössler (x,y; x,z; y,z) cases, although wPLI performed significantly better than wSMI in the intra-hemispheric case of Rössler (x,y). The evaluation of accuracy levels as a function of SNR showed that wSMI tended to perform better than wPLI for intermediate/low SNRs (< 0.45) in the Rössler (y,z) case, while it showed a steep decrease in accuracy at high SNRs (R-R Rössler (x,y) ≥ 0.89; L-R Rössler (x,z) ≥ 0.94; L-R/R-R Rössler (y,z) ≥0.91/0.94). Finally, while wPLI and wSMI showed similar mean accuracy in the Lorenz (x,y) case (with wPLI performing relatively better for SNRs in the range 0.08-0.16), only wSMI was able to detect interactions based on Lorenz (x,z) and Lorenz (y,z) dynamics.

**Figure 5.**
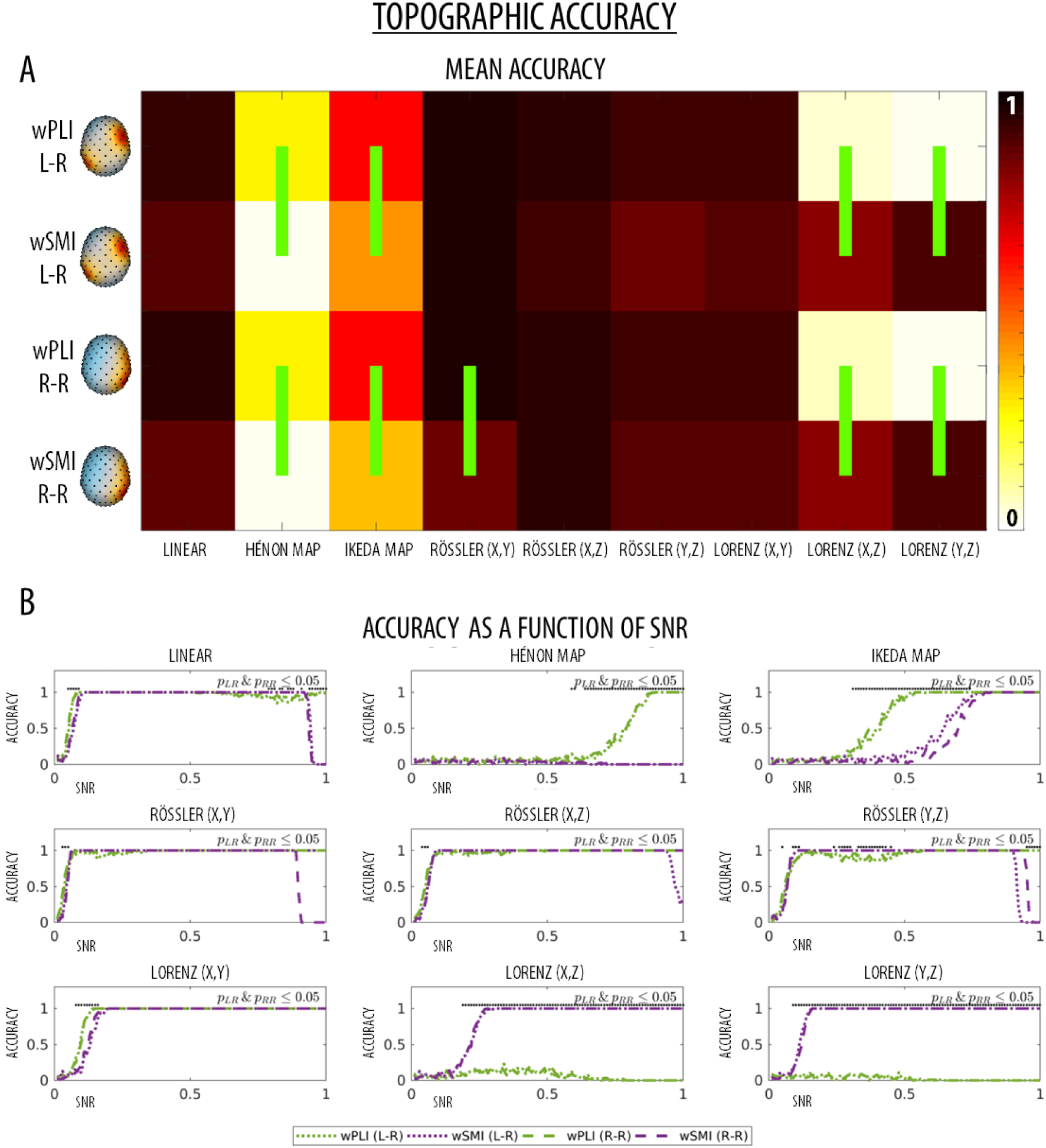
**A)** Mean topographic detection accuracy for all nine different relationships between the chosen source location pairings (L = left IPL to right MFG; R = right IPL to right MFG). The green vertical lines mark significant differences between wPLI and wSMI (permutation tests, p < 0.05) for each type of interaction, pairing of source locations and SNR. **B)** Topographic detection accuracy for all nine different relationships between the chosen source location pairings as a function of SNRs (L = left IPL to right MFG; R = right IPL to right MFG). Black dots at the top of each graph mark significant accuracy differences between wPLI and wSMI for each specific SNR that were observed for both intra and inter-hemispheric conditions.

### Experimental data in wakefulness and sleep

In wakefulness, both wPLI and wSMI revealed significant levels of connectivity in all tested electrodes (p<0.05, cluster-corrected), relative to values observed in time-point shuffled data (Figures 6,7A; 0.5-12 Hz frequency range). In particular, for both measures the highest connectivity values were observed in posterior (occipital, parietal) areas. However, in N3-sleep the two methods provided different results: wPLI revealed diffuse high connectivity values peaking in frontal areas, while wSMI showed reduced connectivity values especially in posterior and lateral electrodes (Figures 7, 7B). In line with these observations, the direct contrast between wakefulness and N3-sleep also revealed distinct changes based on wPLI and wSMI (Figures 6, 7C). Specifically, while wSMI connectivity was significantly higher for wakefulness as compared to N3-sleep in all areas, there were no statistically significant differences in wPLI between these two states of vigilance.

**Figure 6.**
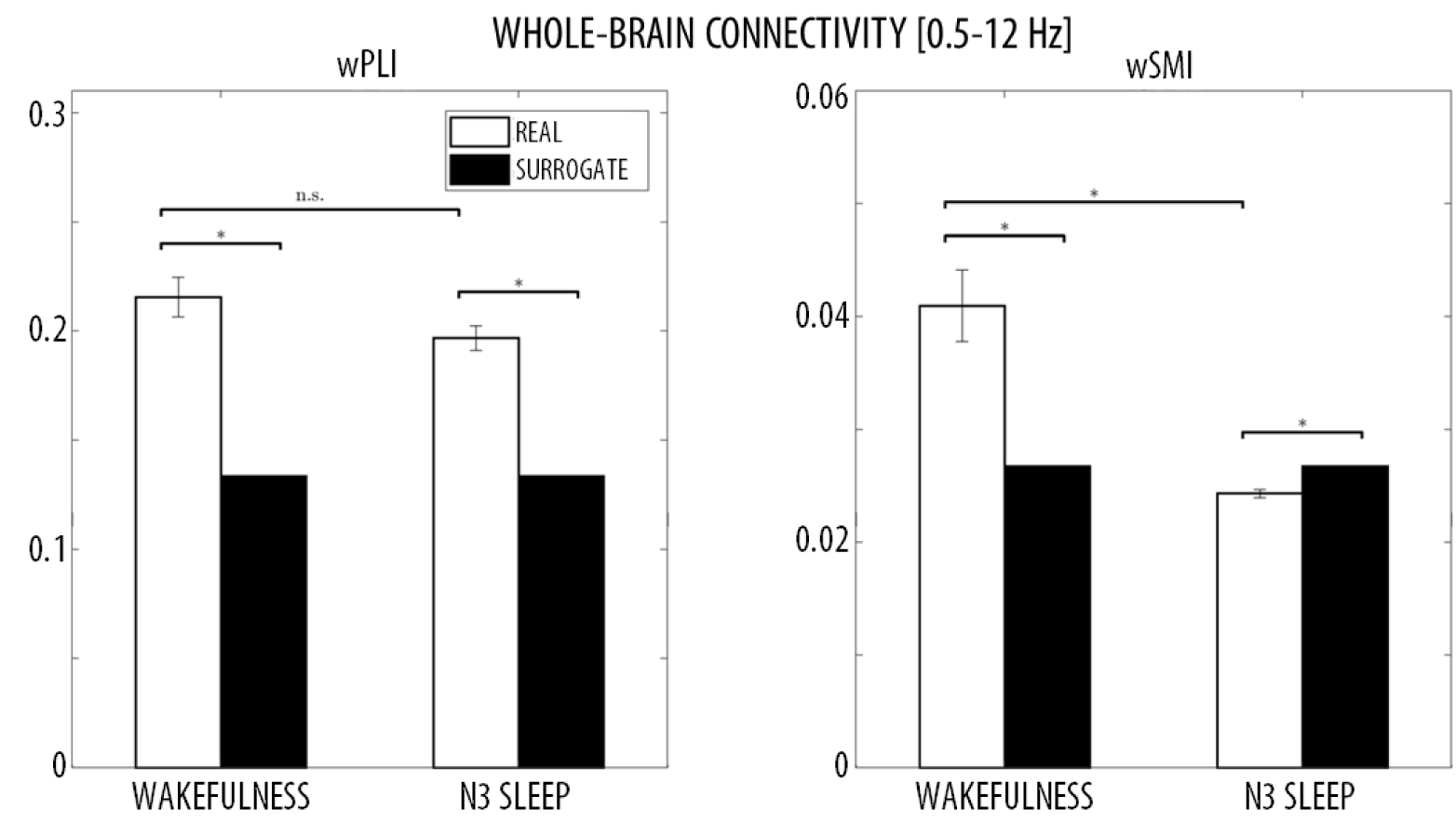
Whole-brain wPLI (left) and wSMI (right) connectivity in wakefulness and sleep (0.5-12 Hz). Paired comparisons were performed between median whole-brain connectivity in wakefulness and N3-sleep, as well as between experimental and surrogate data. * marks p< 0.05 (non-parametric permutation test, N=1000).

**Figure 7.**
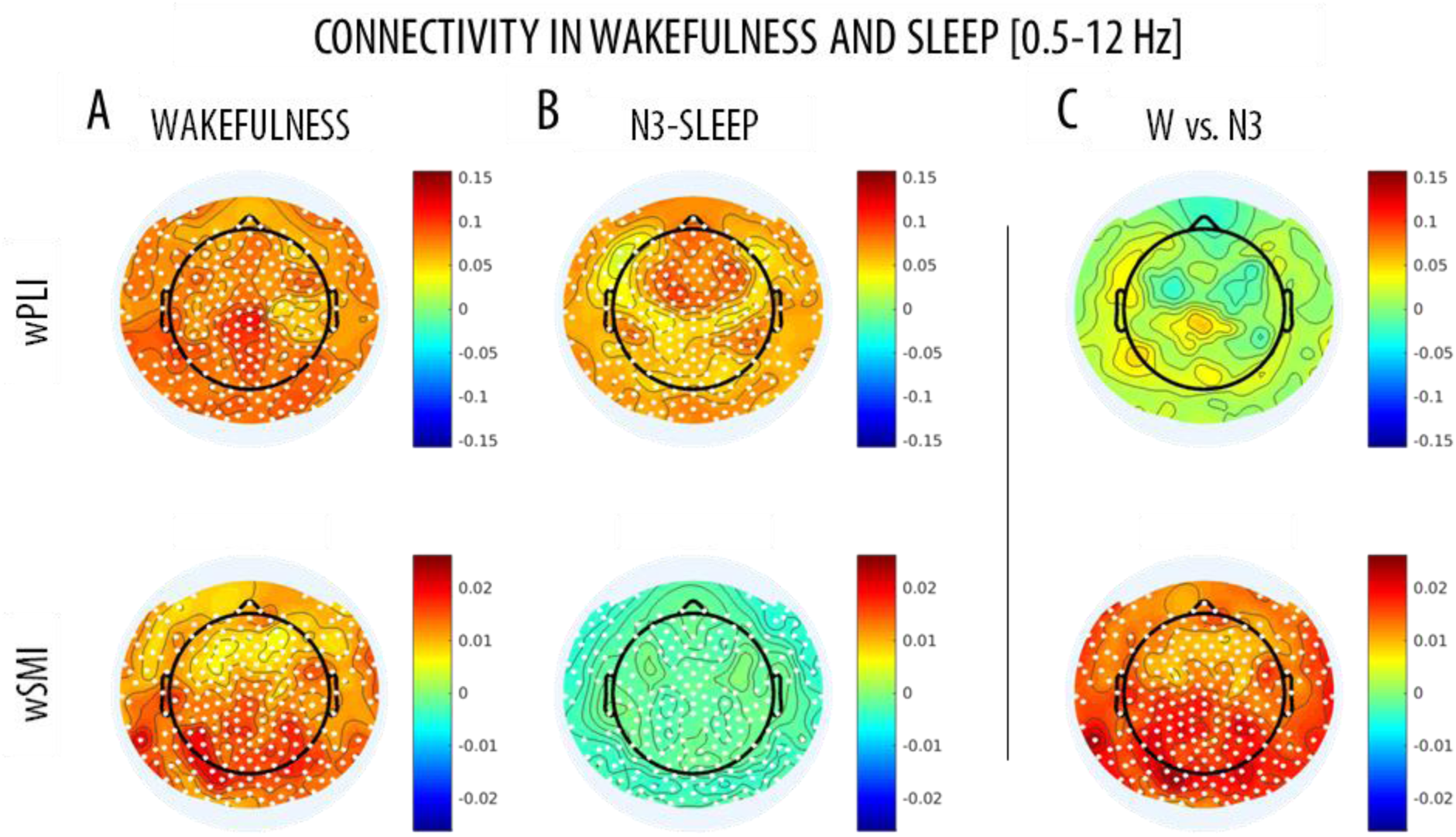
Topographic wPLI (left) and wSMI (right) connectivity in wakefulness and sleep (0.5-12 Hz). Paired comparisons were performed between A) wakefulness and shuffled surrogate data and B) N3-sleep and shuffled surrogate data and C) wakefulness and N3-sleep, for wPLI (top row) and wSMI (bottom row). White dots mark significant effects (cluster-based non-parametric permutation test, p<0.05). Colorbars show the median connectivity difference between conditions, so that the red color marks higher values in real vs. surrogate data for panels A and B. In panel C, the red color indicates higher values in wakefulness, while the blue color indicates higher values in sleep.

Further analyses focusing on classical frequency bands (delta: 0.5-4Hz, theta: 4-8Hz, alpha: 8-12Hz), showed that both wPLI and wSMI were higher in wakefulness than in sleep within the alpha-band (Figures 8-9). However, wPLI was also lower in wakefulness relative to N3 in the delta-band (no differences in theta), while wSMI was lower for wakefulness in the theta-band (no differences in delta).

**Figure 8.**
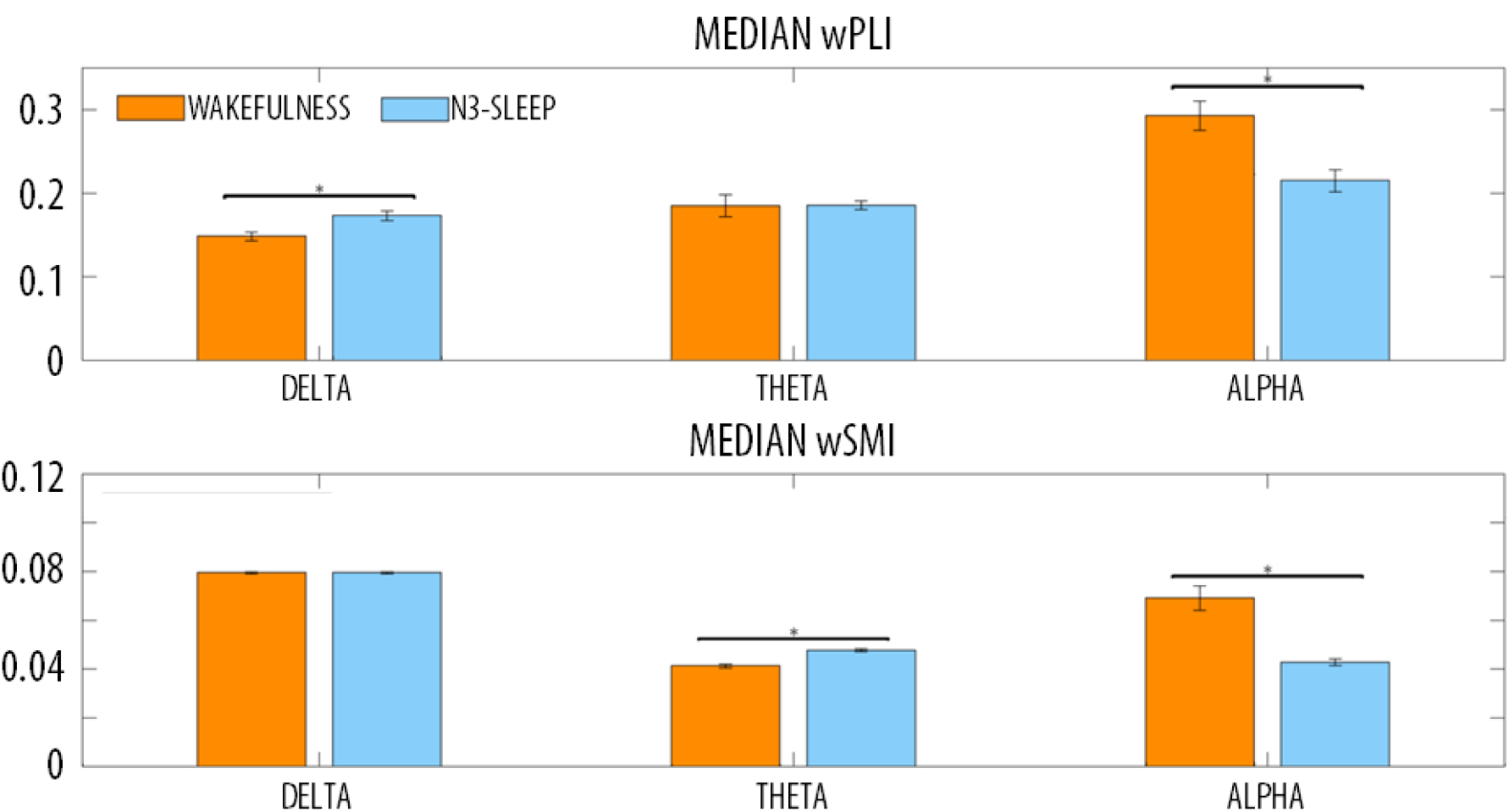
Whole-brain, median wPLI and wSMI in wakefulness (W) and N3-sleep in different frequency bands (delta: 0.5-4Hz, theta: 4-8Hz, alpha: 8-12Hz). Error bars show the standard error of the mean. Horizontal bars and * mark significant differences between conditions (p < 0.05).

**Figure 9.**
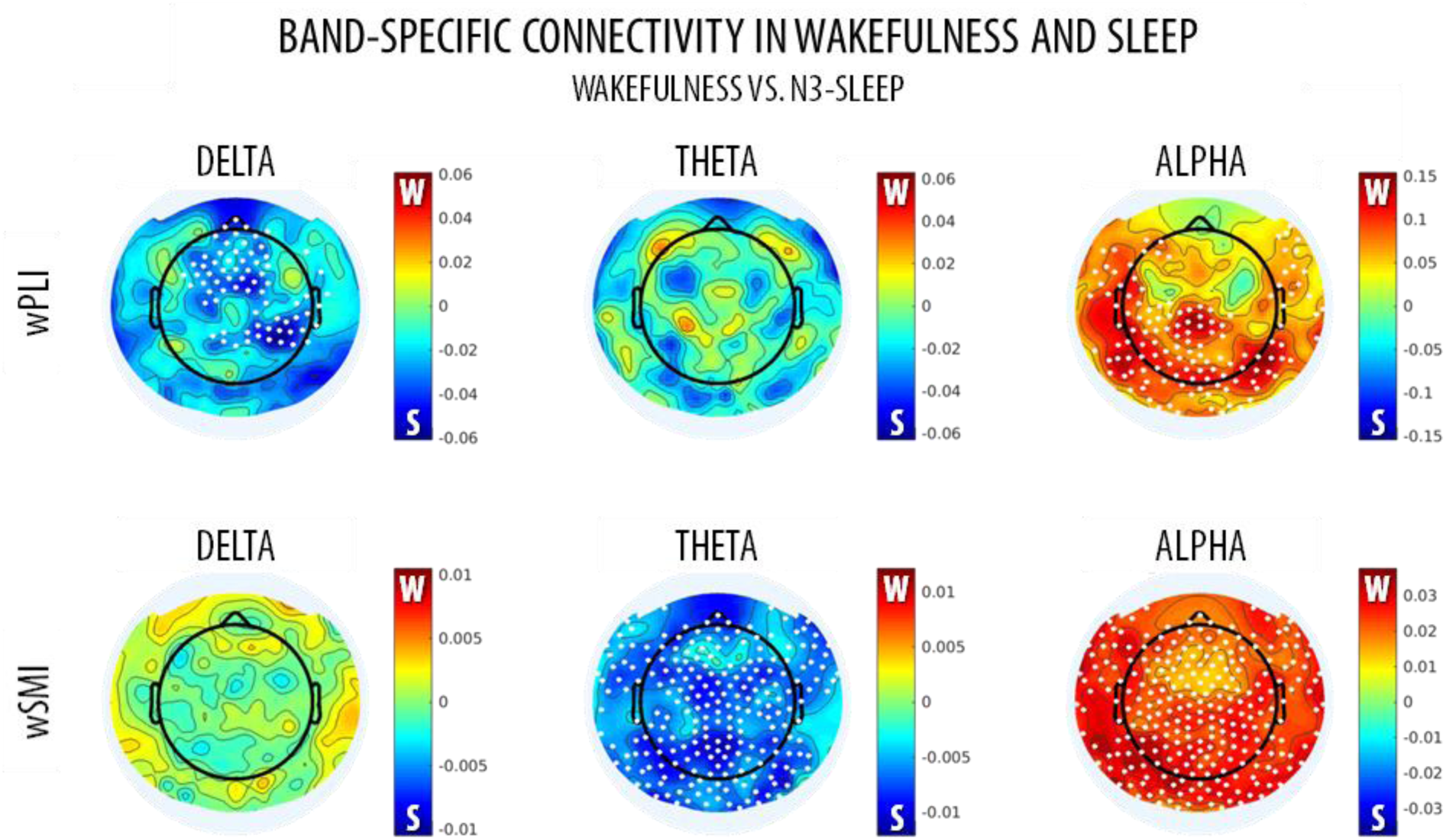
Topographic wPLI and wSMI connectivity in wakefulness (W) and N3-sleep in different frequency bands (delta: 0.5-4Hz, theta: 4-8Hz, alpha: 8-12Hz). Colorbars show the median connectivity differences between wakefulness and N3-sleep. The red color marks higher values in wakefulness, while the blue color indicates higher values in sleep. White dots mark significant differences between conditions (cluster-based non-parametric permutation test, p<0.05).

## Discussion

The wPLI [1] and the wSMI [4] are two robust functional connectivity approaches increasingly applied to M/EEG data, because of their relative immunity to volume conduction effects [5]–[12], [14], [20]. Here we set out to investigate whether the two methods are able to capture overlapping or complementary information regarding variations in brain inter-regional interactions. By combining analyses on simulated hd-EEG data and real hd-EEG recordings collected in different states of vigilance, we demonstrated that wPLI has an optimal sensitivity for interaction dynamics presenting a mixture of linear and nonlinear components, whereas wSMI has higher sensitivity to predominantly nonlinear dynamics. Given that the brain is a highly complex system typically characterised by both linear and nonlinear interaction dynamics [58], it may be better described through the combined use of different measures [26]. Consistent with this view, our results suggest that the conjoint use of wPLI and wSMI may allow researchers to measure complementary information about FC interactions, and thus to better describe relative changes associated with distinct behavioural states.

### Performance of wPLI and wSMI in simulated data

The *Berlin Brain Connectivity Benchmark* (BBCB) framework [29] was adapted and employed to generate hd-EEG recordings in sensor-space. This framework allowed us to model different interaction dynamics between two cortical sources, noise with temporal and spatial structure as well as source mixing due to volume conduction, in a highly realistic electromagnetic volume conductor (head) model. In particular, we generated interaction dynamics with different degrees and types of nonlinearity, from linear to exclusively nonlinear, and specifically tested the sensitivity of wPLI and wSMI at detecting these inter-regional dependencies. Of note, for each of the tested dynamics, we also tested two different source locations (intra and inter-hemispheric interactions) and different signal-to-noise ratios (SNR). Our results showed that the phase-based measure wPLI performs generally better at detecting inter-regional couplings presenting both linear and nonlinear components. Only in two of the more complex nonlinear coupling cases (Lorenz (x,z) and Lorenz (y,z)), characterised by non-significant cross-correlation values (see Figures 4 and 5), wPLI had a very low accuracy. Contrarily, the information-theoretic measure wSMI had a significantly higher accuracy for these two interaction dynamics, but performed significantly worse for the Hénon and Ikeda-based couplings. In particular, the very low accuracy reached by wSMI in the Hénon case may be explained by the high similarity of the source time-series (one time-series represents the scaled and lagged copy of the other one), which may resemble typical results of spurious correlation due to volume conduction. In this respect, the weighting approach applied to wSMI, which actually entirely removes, rather than modulates, the contribution of almost-synchronous variations in EEG signals, may have led to very low connectivity values.

With few exceptions, the accuracies of wPLI and wSMI were very similar for intra and inter-hemispheric interactions, and the detection accuracy of both methods tended to increase with an increase in SNR. Of note, however, the spatial (topographic) accuracy of wSMI (but not the whole-brain accuracy based on median global connectivity) showed instead a decrease at high SNRs for linear and Rössler interactions. This accuracy reduction may be related to an increase in the spatial spreading of the source signals to more distant scalp electrodes with increasing SNRs, which may have led a greater proportion of electrodes to detect the underlying functional coupling (loss of spatial resolution). Moreover, at high SNRs a relative ‘cross-contamination’ may be expected to occur between the two electrodes spatially closest to the interacting sources. In particular, the activity of one source may be ‘volume-conducted’ to the electrode closest to the other source (and vice-versa). Due to the particular weighting approach used for wSMI, the increased similarity between the signals of these particular channels may limit the maximum attainable connectivity strength, thus reducing the relative difference with respect to all other electrode pairings. On the other hand, such effects of volume conduction at high SNRs can be expected to have had only a marginal impact on (or even to improve) the estimation of whole-brain accuracy with respect to null-datasets generated from point or phase-shuffled time-series. The relative difference in the effects of volume conduction for topographic and whole-brain detection accuracy may also contribute to explain the reduced intra-hemispheric (vs. inter-hemispheric) wSMI topographic accuracy observed in the Rössler (x,y) case.

Overall, our results demonstrated that wPLI, as a measure of phase synchronization, performs generally better at detecting functional couplings presenting a mixture of linear and nonlinear dynamics, whereas wSMI, fundamentally rooted in mutual information, has higher sensitivity for exclusively nonlinear dynamics, such as Lorenz (x,z) and Lorenz (y,z) dynamics. Importantly, present results also demonstrated that both wPLI and wSMI are characterised by a high spatial (topographic) accuracy, thus supporting their use in graph theoretical analysis.

### Performance of wPLI and wSMI in distinct states of vigilance

To evaluate whether the results we obtained from simulated EEG data are relevant to the analysis of real experimental data, we tested and compared the performance of the two connectivity measures in hd-EEG recordings collected in humans in different states of vigilance. In fact, both wPLI and wSMI have been previously shown to successfully identify relative variations in brain FC associated with different degrees of consciousness under anesthesia or following severe brain injury [4], [10]–[13], [23]. Based on these premises, here we asked whether the two methods may identify similar or distinct changes associated with variations in the level of consciousness of healthy subjects from wakefulness to deep NREM-sleep (N3). In humans, N3-sleep is characterised by the occurrence of large and diffuse EEG slow waves (0.5-4 Hz), by relative sensory disconnection [59] and by a low probability of having any conscious experiences (dreams) [60]. It has been suggested that slow waves, representing the alternation of neuronal silence (*off-period*) and firing (*on-period*), and occurring out-of-phase in different cortical areas, may contribute to the fading of consciousness through the interruption of causal interactions between distant brain regions [61]–[64].

Here we showed that N3-sleep is associated with a significant and diffuse decrease in wSMI connectivity within the 0.5-12 Hz frequency range. Such difference appeared particularly prominent in posterior brain areas. In contrast, we observed no significant differences between wakefulness and N3-sleep in broadband (0.5-12Hz) wPLI connectivity. A band-limited analysis revealed that changes in wSMI were mainly driven by an overall decrease in *alpha* (8-12 Hz) connectivity in N3 relative to wakefulness. Of note, *alpha*-band wPLI connectivity also showed a similar, but more localized, decrease during N3-sleep, especially in posterior areas. These results are in line with previous work showing that the transition into unconsciousness due to sedation or physiological sleep (stage N1/N2) is associated with a decrease in alpha wPLI-connectivity [12], [13], [23], [65], [66], especially in posterior regions [12], [13] and for posterior-anterior interactions [23], [65], [66]. Moreover, they are consistent with evidence indicating that relative to healthy individuals, patients with unresponsive wakefulness syndrome (UWS) or in a minimally conscious state (MCS) display a connectivity decrease that mainly affects posterior areas or posterior-anterior interactions [4], [10], [11], [24]. Similarly, alpha-band wSMI has been found to be lower in UWS as compared to MCS patients [11]. Therefore, our findings indicate that both wPLI and wSMI may be suited to capture variations in*alpha*-connectivity associated to relative changes in vigilance and/or responsiveness to the environment. However, wPLI and wSMI also identified distinct variations within the other frequency bands. In fact, wSMI identified a diffuse increase in *theta* (4-8 Hz; but not*delta*) connectivity during sleep, while wPLI revealed a relative increase in *delta* (0.5-4 Hz; but not *theta*) connectivity. Importantly, the change in *delta*-wPLI is consistent with the presence of traveling slow waves during sleep [67] as well as with a recent similar observation of increased parietal and parieto-frontal *delta*-wPLI connectivity during propofol sedation [13] and midazolam-based anesthesia [12]. Moreover, wPLI in the *delta*/*theta*-band has been shown to be increased in patients with disorders of consciousness (UWS, MCS), relative to healthy awake subjects [10].

In summary, the analysis of wPLI and wSMI-based connectivity in different states of vigilance confirmed our findings in simulated data, indicating that the two methods are sensitive to distinct brain dynamics. While an in-depth characterization of the differences in FC between wakefulness and sleep was beyond the scope of the present work, our results also suggest that wakefulness may be characterised by a mixture of ‘simple’ (i.e., mainly linear; better described by wPLI) and more complex (i.e., mainly nonlinear) interactions (better described by wSMI) in the alpha range, while sleep may be dominated by ‘simpler’*delta*-band connectivity (better captured by wPLI), likely reflecting the occurrence of traveling slow waves. This interpretation is in line with previous observation indicating that N3 is associated with lower complexity or entropy [58], [68] as compared to wakefulness.

## Conclusions

Our study demonstrates that wPLI and wSMI connectivity metrics provide distinct but complementary information about inter-regional interactions and indicate that the combined use of these two methods may provide a better and more complete characterization of brain functional dynamics within and across distinct behavioural states. In particular, we showed that while wPLI displays an optimal sensitivity for interaction dynamics with linear and nonlinear components, wSMI has a higher sensitivity for predominantly nonlinear dynamics. We also showed that this finding may have significant implications for the analysis of functional connectivity in states of vigilance associated with different levels of consciousness. In light of recent evidence indicating that the independent application of wPLI and wSMI connectivity metrics may allow to identify changes in brain connectivity associated with variations in the level of consciousness, our results point to their possible combined use as a powerful tool to increase their accuracy and predictive value. Nonetheless, our findings may also have more general implications for the study of functional connectivity in a wide variety of behavioural conditions, characterised by distinct underlying brain dynamics.

## Acknowledgments

The authors thank Tristan Bekinschtein for support during the initial planning of the study and Andrea Leo for assistance in the use of the PALM software. This work was supported by a research grant of the Italian Ministry of Health (Ricerca Finalizzata 2011-2012, GR-2011-02347383; to E.R.), the Swiss National Science Foundation (Ambizione Grant PZ00P3_173955), the Divesa Foundation Switzerland, the Pierre-Mercier Foundation for Science, the Bourse Pro-Femme of the University of Lausanne (to F.S.), and a Research Support Grant of the University of Lausanne (to F.S. and G.B.).

## Author contributions (CRediT taxonomy)

Conceptualization, L.S.I, A.C.J., S.C., G.B.; Methodology, L.S.I., M.B., L.C., A.C.J., S.C., G.B.; Investigation, L.S.I., M.B., G.B., F.S.; Formal Analysis, L.S.I.; Visualization, L.S.I., G.B.; Resources, F.S., E.R., P.P.; Supervision, M.B., E.R., P.P, S.C., G.B.; Writing – Original Draft, L.S.I., M.B., S.C., G.B.; Writing – Review & Editing, L.S.I., M.B., L.C., E.R., F.S., P.P., S.C., G.B.

